# Same bi-directional modulation of contrast appearance for voluntary presaccadic attention and involuntary exogenous attention

**DOI:** 10.1101/2024.10.03.616520

**Authors:** Tianyu Zhang, Yongchun Cai

**Author notes:** Correspondence to: Yongchun Cai, Department of Psychology and Behavioral Sciences Zhejiang University, Hangzhou, China.

## Abstract

Different types of attention alter subjective visual perception in fundamentally distinct ways. Previous studies have focused on covert attention without concurrent eye movements, showing that covert exogenous (involuntary) attention enhances contrast appearance of low-contrast stimuli while diminishing that of high-contrast stimuli, whereas covert endogenous (voluntary) attention uniformly enhances contrast appearance. However, the attentional effect preceding saccadic eye movements, a critical component of natural vision, remain understudied. Here, we found that when participants voluntarily initiated saccades, presaccadic attention enhanced the appearance of low-contrast stimuli while attenuating the appearance of high-contrast stimuli (Experiment 1, *N* = 29 adults). This pattern is surprisingly similar to exogenous attention but distinct from endogenous attention. Notably, the presaccadic attentional attenuation effect accumulated gradually during saccade preparation and remained positively correlated with the exogenous attentional effect (Experiment 2, *N* = 24 adults). These findings suggest a shared mechanism between presaccadic and exogenous attention in shaping visual perception.

## Introduction

We are flooded with a vast amount of visual information every time we open our eyes, yet it seems effortless to understand what we see. This can be largely owed to “attention” – a cognitive process that filters relevant information out of irrelevant background to prioritize information processing (Carrasco, 2011; Treue, 2001). The allocation of attention toward a target stimulus is typically achieved through rapid, large eye movements (i.e., saccades) that position the target at the fovea, where the visual acuity is highest (Henderson, 2003). Interestingly, attention shifts toward the target location even before saccade onset, a phenomenon known as “presaccadic attention” (H.-H. Li, Pan, et al., 2021). Presaccadic attention usually enhances visual performance by improving visual sensitivity (H.-H. Li et al., 2016; Rolfs & Carrasco, 2012) and acuity (Kwak et al., 2023), and it modulates the encoding of basic visual features (e.g., spatial frequency and orientation) at the pre-saccadic location (H.-H. Li et al., 2016). In contrast, it imposes perceptual costs for unattended locations (Hanning et al., 2023; H.-H. Li, Pan, et al., 2021). Moreover, presaccadic attention even alters visual appearance, affecting how we subjectively perceive the world. Notably, it enhances contrast appearance of low-contrast stimuli, making them stand out more prominently from the background (Rolfs & Carrasco, 2012).

Deployment of attention may not necessarily accompany saccadic eye movement. Attentional shifts without eye movement are termed covert attention, which constitutes a significant portion of the attention literature (Carrasco, 2011; Posner, 1980; Rizzolatti et al., 1987). Covert attention can be divided into two categories: exogenous attention (involuntary, transient, stimulus-driven, and peaking at ∼100 ms) and endogenous attention (voluntary, sustained, goal-driven, and taking at least 300 ms to deploy) (Cheal & Lyon, 1991; Corbetta & Shulman, 2002; Moore & Zirnsak, 2017; Nakayama & Mackeben, 1989). Both exogenous and endogenous attention can boost perception and performance, via enhancing visual sensitivity, increasing spatial resolution and speeding response to target detection and discrimination (for review, see Carrasco, 2011). However, ample evidence from psychophysical, neurophysiological and neuroimaging studies has demonstrated that these two forms of covert attention exert influence on visual perception in different ways (for reviews, see Carrasco, 2011; Li, Hanning, et al., 2021). For example, exogenous attention consistently increases spatial resolution (Carrasco et al., 2006), whereas endogenous attention can either enhance or decrease spatial resolution, depending on task demands (Barbot & Carrasco, 2017; Jigo & Carrasco, 2018; Yeshurun et al., 2008). Exogenous attention increases contrast sensitivity through mechanisms of signal enhancement (Cameron et al., 2002; Carrasco et al., 2000) and external noise reduction (Lu & Dosher, 1998), whereas endogenous attention enhances contrast sensitivity solely by reducing external noise (Lu & Dosher, 2000). Recent studies indicate that the two types of cover attention influence contrast appearance differently: both exogenous and endogenous attention enhance contrast appearance for low-contrast stimuli (Carrasco et al., 2004; Störmer et al., 2009), but for exogenous attention, the enhancement effect turns into an attenuation effect as stimulus contrast increases (Itthipuripat et al., 2019; Pan et al., 2023; Pan & Cai, 2022; Zhou et al., 2018). Conversely, endogenous attention continues to enhance contrast appearance at higher contrast levels (Luo et al., 2024). These findings underscore that covert exogenous and endogenous attention have fundamentally different underlying neurocognitive mechanisms for modulating visual perception.

Presaccadic attention is closely related with, though dissociable from, both covert exogenous and endogenous attention at behavioral and neural level (H.-H. Li, Hanning, et al., 2021; Moore & Zirnsak, 2017; Smith & Schenk, 2012). The premotor theory of attention even posits covert attention as subsidiary of prepared but unexecuted saccades, which would result in presaccadic attention if prepared saccades were executed (Rizzolatti et al., 1987; Smith & Schenk, 2012). Despite the close linkage, few studies have directly compared the perceptual effects of presaccadic attention with those of covert exogenous attention and endogenous attention. Specifically, when people voluntarily initiate saccades toward target locations, does presaccadic attention modulate visual perception in ways more similar to exogenous or endogenous attention? Intuitively, goal-directed presaccadic attention should resemble covert endogenous attention. This intuition is implied by various computational models from reading to visual search (Einhäuser et al., 2008; Engbert et al., 2005; X. Li & Pollatsek, 2020; Rao et al., 2002; Reichle et al., 2006; Snell et al., 2018), and is strengthened by evidence that the frontal eye field (FEF) serves as a shared neural origin for both endogenous attention and goal-directed saccades (Clark et al., 2015; Fernández et al., 2023; Moore & Fallah, 2004).

However, recent evidence suggests that presaccadic attention, even preceding voluntary saccades, more closely mirrors covert exogenous attention in modulating visual perception. For instance, exogenous attention is impaired for targets outside the oculomotor range (Casteau & Smith, 2020; Smith et al., 2012; but see Hanning et al., 2019) or in patients unable to move their eyes (Gabay et al., 2010; Sereno et al., 2006; Smith et al., 2004), whereas endogenous attention remains unaffected in these cases (but see Craighero et al., 2001). Additionally, both presaccadic and exogenous attention arbitrarily enhance high spatial frequency processing, even if this impairs task performance (Jigo et al., 2021; H.-H. Li et al., 2019); conversely, endogenous attention flexibly adjusts high spatial frequency processing to optimize task performance (Barbot & Carrasco, 2017). TMS studies further highlight a causal role of primary visual cortex (V1) in mediating effects of presaccadic and exogenous attention, but not endogenous attention (Fernández et al., 2023; Fernández & Carrasco, 2020; Hanning et al., 2023). Together, these findings suggest a potential similarity between voluntary presaccadic attention covert exogenous attention in shaping visual perception, though direct investigations are still lacking.

Here, we aimed to explore the potential link between presaccadic and covert attention by examining their perceptual consequences. We investigated the voluntary presaccadic attentional effect on contrast appearance at low and high contrast levels and compared it to the effects of covert attention. If presaccadic attention were more tightly linked to covert endogenous attention, we expected a uniform attentional enhancement effect across contrast levels. However, if presaccadic attention were more tightly linked to covert exogenous attention, we anticipated a bi-directional attentional modulation, with enhancement for low-contrast and attenuation for high-contrast stimulus. Our findings confirmed the latter: voluntary presaccadic attention alters contrast appearance bi-directionally, identical with exogenous attention but contrary to endogenous attention. Notably, the effects of presaccadic attention gradually built up during saccade preparation but persistently maintained a positive correlation with effects of exogenous attention. These results suggest that the two types of attention engage a common mechanism in modulation of subjective visual perception, despite being behaviorally allocated in fundamentally different ways.

## Method

### Participants

We recruited a total of 54 college students from Zhejiang University in China as participants. All had normal or corrected-to-normal vision and no history of mental diseases. With the exception of the first author, ZTY, all participants were naïve to the purpose of the experiment. Informed consent was obtained from each participant, who received either monetary compensation or course credits for their involvement. The Research Ethics Board of Zhejiang University approved the experimental procedures and protocols.

Eighteen participants attended Experiment 1a (the low-contrast stimulus condition) and 18 attended Experiment 1b (the high-contrast stimulus condition), with seven participating in both. The sample size was determined based on a recent study by Zhou et al. (2018), who employed an equality judgment task to explore how attention influences contrast perception. In their study, the average effect size (*d*_z_) under high-contrast conditions was 1.7. Consequently, a sample size of 7 was required to achieve a statistical power of 0.95, while maintaining a significance level of 0.05 (two-tailed; computed with G*Power 3.1, Faul et al., 2009). We have increased the sample size to adequately detect the potential effects after excluding participants with poor fit of psychophysical curve. Data from 4 participants in experiment 1a and 3 in Experiment 1b were excluded from further analyses due to poor psychometric curve fits (*R*^2^ < .5). This left a total of 14 participants for experiment 1a (5 females, aged 18-25) and 15 participants for experiment 1b (6 females, aged 18-27).

Thirty-one participants attended Experiment 2, with six having also participated in Experiment 1. Seven participants were excluded due to poor psychometric curve fits (*R*^2^ < .5), leaving 24 valid participants (12 females, aged 18-27) for Experiment 2.

### Apparatus

All stimuli were generated using Psychophysics Toolbox (Brainard, 1997; Pelli, 1997) and Eyelink Toolbox extensions (Cornelissen et al., 2002) in MATLAB (MathWorks, Natick, MA, USA), displayed on a gamma corrected CRT monitor (19′′ Dell P992; 1024 × 768 resolution; 100 Hz refresh rate). Eye positions were recorded by the Eyelink 1000 Plus eye tracker (SR Research, Osgoode, Ontario, Canada) at a sampling rate of 1 kHz. The viewing distance was 70 cm from the screen with the head stabilized by a chinrest.

### Stimuli

Sequences of stimuli during a trial for Experiments 1 and 2 are illustrated in Fig. 1. The fixation stimulus (a dot with diameter of 0.3° in a black circle with diameter of 0.7°) was present at the center of the screen throughout the experiment against a grey background (38.1cd/m^2^). Placeholders measuring 4°×4° visual angle, consisted of 4 dots (Exp. 1: 0.2° × 0.2° black dots; Exp. 2: 0.15° × 0.15° grey dots), were located on the left and right side of the fixation (eccentricity 8°) and were presented in all conditions of Experiment 1 and the saccade condition in Experiment 2.

**Fig. 1.**
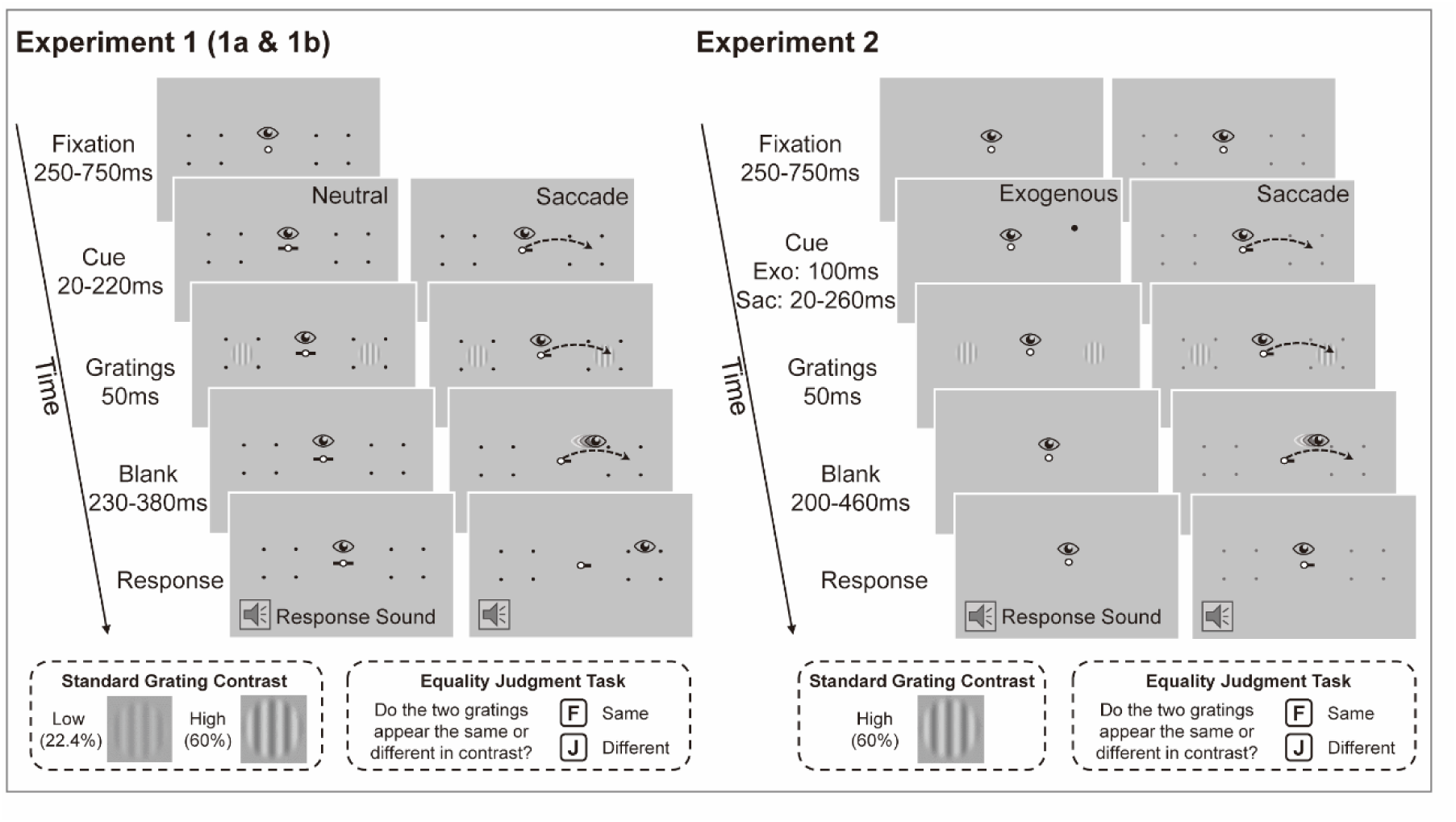
Schematic representation of stimuli and procedures in Experiment 1 (left) and Experiment 2 (right). In both experiments, participants judged whether the two target gratings had the same or different contrast. Both Experiments 1a (low-contrast stimuli condition) and 1b (high-contrast stimuli condition) included saccade blocks, where participants made saccades toward the cued placeholders, and neutral blocks, where participants maintained central fixation. Experiment 2 included saccade and exogenous blocks and only tested the high-contrast stimuli condition. The procedure for the saccade blocks in Experiment 2 was identical to that in Experiment 1. In exogenous blocks, participants were required to maintain central fixation. The icons of eyes, arrows and trumpets were utilized merely for illustrative purposes and were not shown in experiments.

Target stimuli comprised a pair of vertical sinusoidal gratings (diameter 3°; spatial frequency 1.5 cpd) with borders blurred using a raised cosine function (0.5° width). One grating served as the standard stimulus, with a fixed Michelson contrast of 22.4% in Experiment 1a and 60% in Experiments 1b and 2. The other grating was the test stimulus, with contrast systematically adjusted in 11 log increments (8% to 63% in Experiment 1a; 37% to 97% in Experiments 1b and 2) following an adaptive staircase rule (see details in Pan & Cai, 2022). The two gratings were randomly positioned to the left and right side of the fixation at 8° eccentricity along the horizontal meridian. The phases of the two gratings were identical in each trial but randomized across trials to prevent afterimage.

In the saccade condition of all experiments, a central cue (0.6° length, 0.2° width) pointing either left or right was used to direct saccades. In the neutral condition (Experiment 1), cues pointed to both left and right, requiring maintenance of fixation at the center point. In the exogenous attention condition (Experiment 2), a peripheral black dot (0.7°× 0.7°) positioned 0.9° above the edge of the target grating was used to direct exogenous attention.

### Procedure

The procedures for Experiments 1a and 1b were identical (Fig. 1), except that the grating contrast was low in Experiment 1a and high in Experiment 1b. Experiment 1 consisted of the saccade and neutral conditions. To avoid prolonged saccade reaction times in the saccade condition, the two conditions were blocked (H.-H. Li, Pan, et al., 2021; Rolfs & Carrasco, 2012). Each trial began with participants pressing the space bar, causing a red fixation dot to appear. Participants then had to maintain stable fixation within a 1.5° radius around the fixation point for 200ms. If stable fixation was not achieved within 3 seconds, the trial was aborted, and the eye tracker was recalibrated. Upon successful stable fixation, the red dot turned white, followed by a saccade or neutral cue after a random interval of 250-750 ms. The saccade cue pointed randomly left or right, and participants were instructed to saccade towards the center of the placeholder as quickly and accurately as possible. The neutral cue, pointing in both directions, instructed participants to maintain central fixation throughout the trial. Shortly after the cue onset, a pair of gratings flashed 50 ms. An auditory tone, indicating the start of the response phase, was played either after a successful saccade landing in the saccade task or following a 200 ms delay after the grating disappeared in the neutral task. Participants then reported whether the two gratings had the same or different contrast by pressing the “F” or “J” on the keyboard, respectively. No feedback was given. Participants were informed that their contrast judgment was non-speeded and were encouraged to rest and blink as needed before initiating the next trial. This equality judgment task is believed to minimizes response bias (Schneider & Komlos, 2008) and has been widely used to assess the attentional effect on contrast appearance (Itthipuripat et al., 2019; Luo et al., 2024; Pan et al., 2023; Pan & Cai, 2022; Zhou et al., 2018).

Experiment 2 included both saccade and exogenous attention conditions, organized in a block design. The procedure for the saccade condition was identical to that of the saccade condition in the Experiment 1. For the exogenous attention condition, the procedure was similar to the neutral condition in Experiment 1, except that the neutral center cue was replaced with a brief peripheral dot (50 ms) serving as the exogenous attention cue.

The stimulus onset asynchrony (SOA) between the cue and target gratings was selected to optimize the attentional effect for different types of attention. For the saccade condition, the SOA ranged from 20-220 ms in Experiment 1 and 20-260 ms in Experiment 2. These ranges allowed the presentation of gratings before saccade onset in most saccade trials. The SOA for a specific saccade trial was determined using an adaptive algorithm based on the history of saccadic reaction times to maximize the proportion of trials in which the target gratings disappeared before saccade onset. This algorithm was used due to substantial variability in saccadic reaction time within and between participants (Hanning & Deubel, 2022; Harrison et al., 2013). Specifically, for each saccade trial, we estimated the median saccadic reaction time from preceding valid saccades (see below for the criterion). At the beginning of each block, the median saccadic reaction time was estimated from valid trials of previous blocks, except for the first block, which used data from the practice session. The SOA for a saccade trial was determined by subtracting a random duration (Experiment 1: 100 or 120 ms; Experiment 2: 80, 100, 120, 150, or 170 ms) from the median reaction time. SOAs outside a predefined range (Experiment 1: 20-220 ms; Experiment 2: 20-260 ms) were adjusted to the nearest limit. A wider SOA range was used in Experiment 2 to obtain a more broadly distributed set of presaccadic trials for analyzing the temporal evolution of presaccadic attention. Additionally, in Experiment 2, median saccadic reaction times were separately estimated for trials with different saccade directions, and trials where the stimulus disappeared after saccade onset were redone at the end of the block. These operations ensured that the gratings disappeared before saccade onset in most trials during offline analyses (Experiment 1: 87.3±7.1%; Experiment 2: 93.8±3.8%). For the neutral condition in Experiment 1, the SOA was randomly sampled from a uniform distribution of 20-220 ms to match the SOA in the saccade condition. For the exogenous attention condition in Experiment 2, the SOA was fixed at 100 ms (50 ms cue during plus 50 ms inter-stimulus interval), the point at which exogenous attention typically reaches its peak (Carrasco, 2011).

In Experiment 1, participants completed six saccade and three neutral blocks on two different days. In Experiment 2, participants completed ten saccade and two exogenous attention blocks on two different days. Trials not meeting the online eye movement criteria (see below) were aborted and redone in random order at the end of each block. Each block contained at least 160 trials. Saccade trials outnumbered neutral and exogenous attention trials because we needed to screen for presaccadic trials and discard considerable trials without valid saccades after both online and offline screening. Prior to the formal experiment, participants performed 48 or 96 training trials for contrast judgment with stimulus parameters identical to the neutral condition, except that auditory feedback was provided for incorrect responses.

### Data analysis

#### Online eye movement data analysis and trial exclusion criteria

In all trials, online saccade onset was defined as the time point when gaze position crossed a circular boundary with a radius of 1.5° around the fixation dot. A trial was considered invalid if participants: 1) Blinked at any time before the response sound was played; 2) Looked 1.5° away from the central fixation point before cue onset in the saccade condition, or before the response sound in the neutral and exogenous attention conditions; 3) Failed to initiate a saccade within 480 ms after cue onset in the saccade condition; 4) Landed a saccade more than 2° away from the center of placeholders in saccade trials. These invalid trials were repeated in random order at the end of each block.

#### Offline eye movement data analysis and trial inclusion criteria

In the offline analysis, we utilized the whole eye-position data of each trial to achieve more accurate saccade detection. Using an established algorithm (Engbert & Mergenthaler, 2006; Schweitzer & Rolfs, 2020), we transformed raw eye-position data into two-dimensional velocity distributions. The onset and offset of saccades were identified as the time point when velocities exceeded the median velocity of preceding eye samples by 5 standard deviations for at least 8 ms. Saccades separated by less than 20 ms were merged into a single saccade. Based on the offline-detected saccades, saccadic reaction time was calculated as the duration from cue onset to saccade onset. Landing precision was calculated as the Euclidean deviation between the saccade offset position and the center of placeholders.

In the saccade condition, a trial was considered valid only if the trial had a response saccade leaving from the central fixation (within 1.5° radius) and landing in the placeholder center (within 2.1° radius), with appropriate reaction time between 70-500 ms and without other saccades preceding the response saccade. For both Experiments 1 and 2, saccade trials in which targets presented in the time window 100 - 0 ms before saccade onset were taken into analysis to investigate the presaccadic attention effect (mean trials ± standard deviation in Experiment 1: 280±67 per cued condition; mean trials ± standard deviation in Experiment 2: 405±131 per cued condition).

In addition, for the saccade condition in Experiment 2, we examined the temporal evolution of presaccadic attention by utilizing all valid presaccadic trials. First, for each participant, we divided all valid presaccadic trials into two timebins with equal number of trials (308±49 trials per cued condition, mean ± standard deviation), and assessed the attentional effect within each bin. We calculated the median relative timing between grating offset and saccade onset for each bin as its representative timing and averaged these medians across participants. We below referred to the two timebins as the early (averaged 122±31 ms before saccade onset, mean ± standard deviation) and late stages (averaged 49±18 ms before saccade onset, mean ± standard deviation) of saccade preparation, based on their temporal proximity to saccade onset. Second, we conducted a sliding window analysis to characterize a finer-grained temporal evolution of the attentional effect and its correlation with the effect of exogenous attention. The sliding window size was set to one-third of each participant’s valid presaccadic trials, moving 20 times from the most distant to the most proximal window relative to saccade onset. Representative timing was determined as before. Data from some participants were discarded within certain windows due to poor psychometric curve fitting (on average 1.7 data points were removed per window).

In the neutral (Experiment 1) and exogenous attention (Experiment 2) conditions, trials with saccades before the response cue were excluded. Overall, participants maintained fixations well, with an average of 1.2±2.3 neutral trials and 0.8±3.2 exogenous trials excluded for each participant.

#### Behavioral data analysis

Only trials meeting the offline inclusion criteria described above were included in the behavioral analysis. For each participant, proportions of ‘same’ judgment trials were calculated at each contrast level of test stimuli in each cue condition. The raw contrast levels were converted into logarithmic contrast difference between the test and standard gratings, calculated as Δc = *log_10_*(*c_test_* /*c_standard_*). This value was fitted with a Gaussian function (Pan & Cai, 2022; Schneider & Komlos, 2008) using *fitnlm* function in Matlab. The point of subjective equality (PSE) was defined as the Δc corresponding to the maximum of this bell-shaped function, representing the point at which participants were most likely to perceive the two gratings as having the same contrast. The Δc PSE were subsequently converted back to original contrast scales for statistical analysis. The psychophysical fitting and eye data analyses were conducted in MATLAB using custom scripts and the statistical analyses were conducted using JASP (2023; Version 0.18.1) and R (2023; version 4.3.2). For the ANOVAs, Greenhouse-Geisser-corrected *p* values were reported when there was a violation of the sphericity assumption. Bonferroni correction was used to correct the reported *p* values when multiple *t* tests were conducted.

## Results

### Experiment 1: Presaccadic attention alters appearance bi-directionally

#### Attentional effects on contrast appearance

We measured the effects of voluntary presaccadic attention on contrast appearance in Experiments 1a and 1b for low- and high-contrast stimuli, respectively. Figures 2a and 2d illustrate the probability of participants judging the two gratings as having the same contrast, plotted against the test grating contrast, along with the psychometric curves averaged across participants. The bell-shaped function fitted the behavioral data well (*R^2^* = .85, *SD* = 0.12). A two-way mixed ANOVA on PSEs revealed a significant interaction between the cue and contrast conditions (*F*(1.60, 43.28) = 17.01, *p* < .001, *η*^2^_*p*_ = 0.39), suggesting that presaccadic attention affects contrast appearance differently for low- versus high-contrast stimuli.

**Fig. 2.**
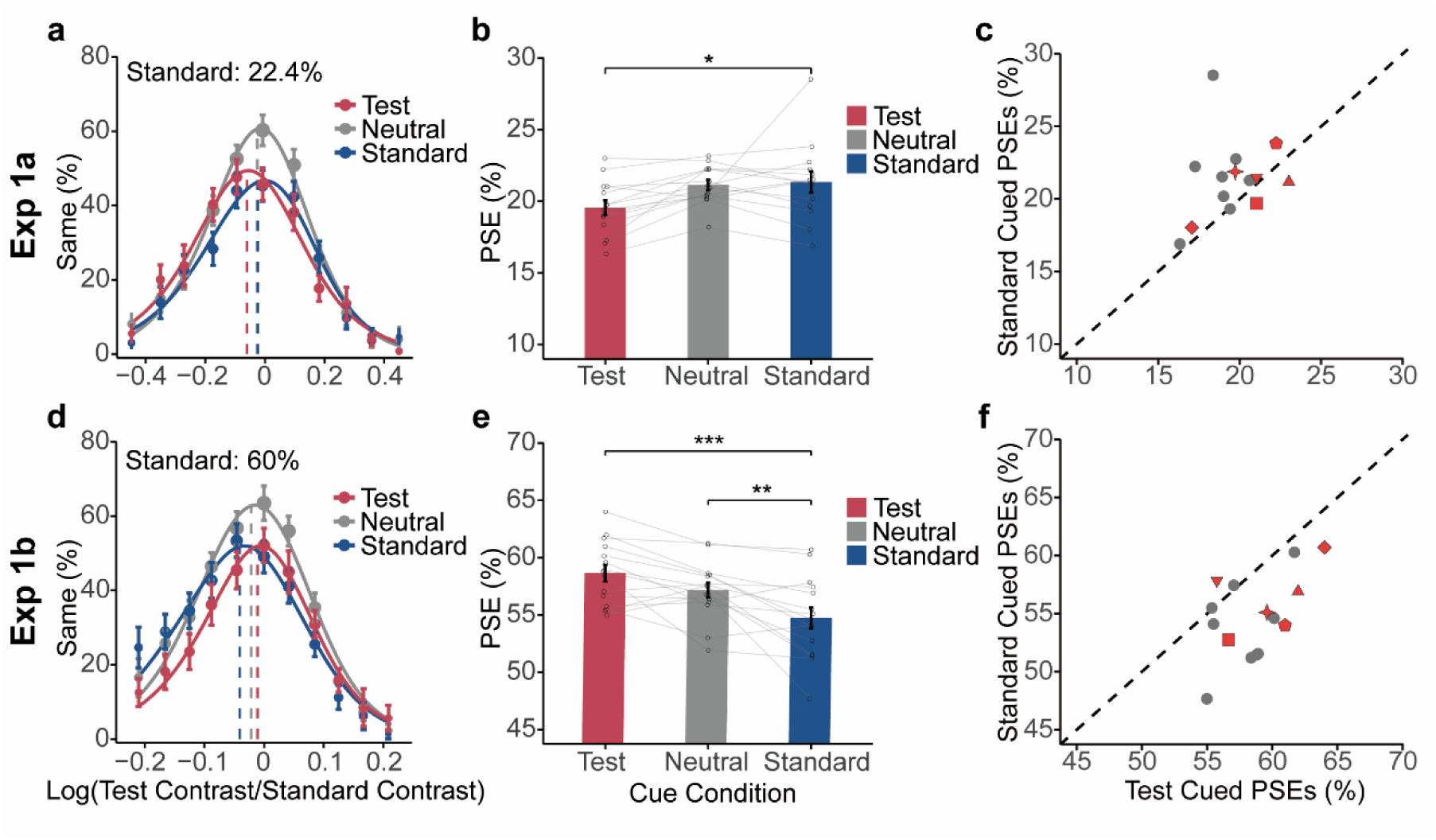
Results of Experiments 1a (low-contrast condition, upper row) and 1b (high-contrast condition, lower row). The left column (a, b) displays psychometric functions, plotting the percentage of ‘same contrast’ responses as a function of the test grating contrast, with each data point size proportional to the number of trials at the corresponding contrast level of the test grating. The middle column (b, e) shows the averaged PSEs at each cued condition, with each data point representing one participant. The right column (c, f) demonstrates individual PSEs in the standard-cued versus the test-cued conditions, where each point denotes one participant. Gray circles represent participants who participated in only one experiment, while red points with unique shapes indicate participants who completed both experiments. The diagonal line signifies equivalent PSEs under test- and standard-cued conditions, suggesting no attentional effect on contrast appearance. Error bars represent ±1 standard errors of the mean. *p < .05. **p < .01. ***p < .001.

To follow up the interaction effect, separate one-way repeated-measures ANOVAs were performed for each contrast condition. Figures 2b and 2e illustrate the group-averaged PSEs under three cue conditions for the low- and high-contrast conditions, respectively. For the low-contrast condition (Experiment 1a), presaccadic attention altered the contrast appearance, *F*(1.36, 17.68) = 4.68, *p =* .035, *η^2^_p_* = 0.27. When presaccadic attention was allocated to the test/standard grating, the curve shifted leftward/rightward compared to the the neutral condition. A post-hoc *t*-test confirmed this pattern, revealing a significantly lower PSE in the test-cued condition than in the standard-cued condition (*t*(13) = 2.78, *p =* .03, *Cohen’s d* = 0.85, 95% CI for the mean difference = [0.14, 3.43]). There was a marginally significant lower PSE in the test-cued condition compared to the neutral condition (*t*(13) = 2.49, *p =* .059, *Cohen’s d* = 0.76, 95% CI for the mean difference = [−0.05, 3.24]), although no significant difference was found between the standard-cued and neutral conditions (*t*(13) = 0.29, *p =* .99, *Cohen’s d* = 0.09, 95% CI for the mean difference = [-1.45, 1.83]). This result indicated that presaccadic attention enhanced the perceived contrast of low-contrast stimuli, replicating previous findings (Rolfs & Carrasco, 2012).

For the high contrast condition (Experiment 1b), presaccadic attention also altered the contrast appearance (*F*(2, 28) = 14.62, *p <* .001, *η^2^_p_* = 0.51). However, the pattern was reversed, as test/standard-cued curves shifted rightward/leftward compared to the neutral condition. A post-hoc *t*-test confirmed this pattern, showing that the PSE in the standard-cued condition was significantly lower compared to the neutral condition (*t*(14) = 3.33, *p =* .007, *Cohen’s d* = 0.82, 95% CI for the mean difference = [−4.29, −0.57]) and the test-cued condition (*t*(14) = 5.36, *p <* .001, *Cohen’s d* = 1.32, 95% CI for the mean difference = [−5.77, −2.05]). However, the PSE difference between the neutral and test-cued conditions was not significant (*t*(14) = 2.03, *p =* .16, *Cohen’s d* = 0.50, 95% CI for the mean difference = [−3.34, 0.38]). This result indicated that presaccadic attention attenuated perceived contrast of high-contrast stimuli. The bi-directional modulation of attention on contrast appearance was also robust at the individual level (Figures 2c and 2f).

#### Possible effects of eye movement

To rule out the possibility that the bi-directional attentional effect observed for low- and high-stimuli was due to differences in saccadic kinematics, we scrutinized eye movement characteristics under both conditions. Our analyses revealed no significant difference in saccadic reaction time (*t*(27) = 0.520, *p =* .61, *Cohen’s d* = 0.19, 95% CI for the mean difference = [−44.81, 26.60]) or landing precision (*t*(27) = 1.05, *p =* .302, *Cohen’s d* = 0.39, 95% CI for the mean difference = [−0.19, 0.06]) across different contrast conditions. The primary factor influencing the magnitude of the presaccadic attention effect is the relative timing between target offset and saccade onset (H.-H. Li, Hanning, et al., 2021). This timing was precisely controlled by an online adaptive algorithm (see Method), showing no significant differences between low-contrast (−52.5 ± 12.0 ms) and high-contrast (−56.5 ± 14.4 ms) conditions (*t*(27) = 0.81, *p =* .43, *Cohen’s d* = 0.30, 95% CI for the mean difference = [−14.15, 6.14]). Therefore, the bi-directional attentional effect cannot be attributed to saccadic kinematic factors.

In summary, Experiment 1 demonstrated that presaccadic attention, intentionally allocated by participants, enhanced the perceived contrast of low-contrast stimuli while diminishing it for high-contrast stimuli. This bi-directional pattern of attentional effects resembles those of covert exogenous (involuntary) attention (Pan et al., 2023; Pan & Cai, 2022; Zhou et al., 2018), but differs from those of covert endogenous (voluntary) attention (Luo et al., 2024). This finding raises an important question: does this similarity between presaccadic attention and exogenous attention suggest common underlying mechanisms, or is it merely a coincidental convergence of the two distinct attention systems? To address this question, we conducted Experiment 2, assessing the correlation between the attentional effects of presaccadic and exogenous attention on high-contrast stimuli using a within-participant design.

### Experiment 2: Correlation of attentional effects by presaccadic and exogenous attention

#### Attentional effects on contrast appearance

Fig. 3a shows the average percentage of “same” responses, plotted as a function of the test contrast. The bell-shaped psychometric functions were averaged across participants under different attention conditions, fitting the behavior data well [pre-saccadic attention: *R*^2^ = .85, *SD* = 0.10; exogenous attention: *R*^2^ = .80, *SD* = 0.14]. To assess the cue effect, separate paired samples *t*-tests were conducted for the presaccadic and exogenous attention conditions. The presaccadic attentional effect shown in Fig. 3 was calculated using trials within 100 ms before saccade onset, consistent with the procedure in Experiment 1. As expected, both types of attention attenuated the perceived contrast of high-contrast stimuli, with a lower averaged PSE in the standard-cued condition than in the test-cued condition (Fig. 3b; presaccadic: *t*(23) = 4.30, *p <* .001, *Cohen’s d* = 0.88, 95% CI for the mean difference = [−5.28, 1.85]; exogenous: *t*(23) = 6.67, *p <* .001, *Cohen’s d* = 1.37, 95% CI for the mean difference = [−8.26, −4.36]). These effects were consistent at the individual level (Fig. 3c). Additionally, a two-way repeated measures ANOVA revealed a significant interaction between the type of attention and cue condition, *F*(1,23) = 9.02, *p =* .006, η*_p_ ^2^* = 0.28, suggesting different magnitudes of attenuation effects for these two types of attention. A follow-up paired samples *t-*test on their cue effects (calculated by subtracting the PSE in the test-cued condition from the PSE in the standard-cued condition) showed that covert exogenous attention had a stronger attenuation effect than presaccadic attention (*t*(23) = 3.02, *p =* .006, *Cohen’s d* = 0.61, 95% CI for the mean difference = [−4.63, −0.85]). These results confirmed the robust attenuation effect of both presaccadic and exogenous attention on contrast appearance of high-contrast stimuli.

**Fig. 3.**
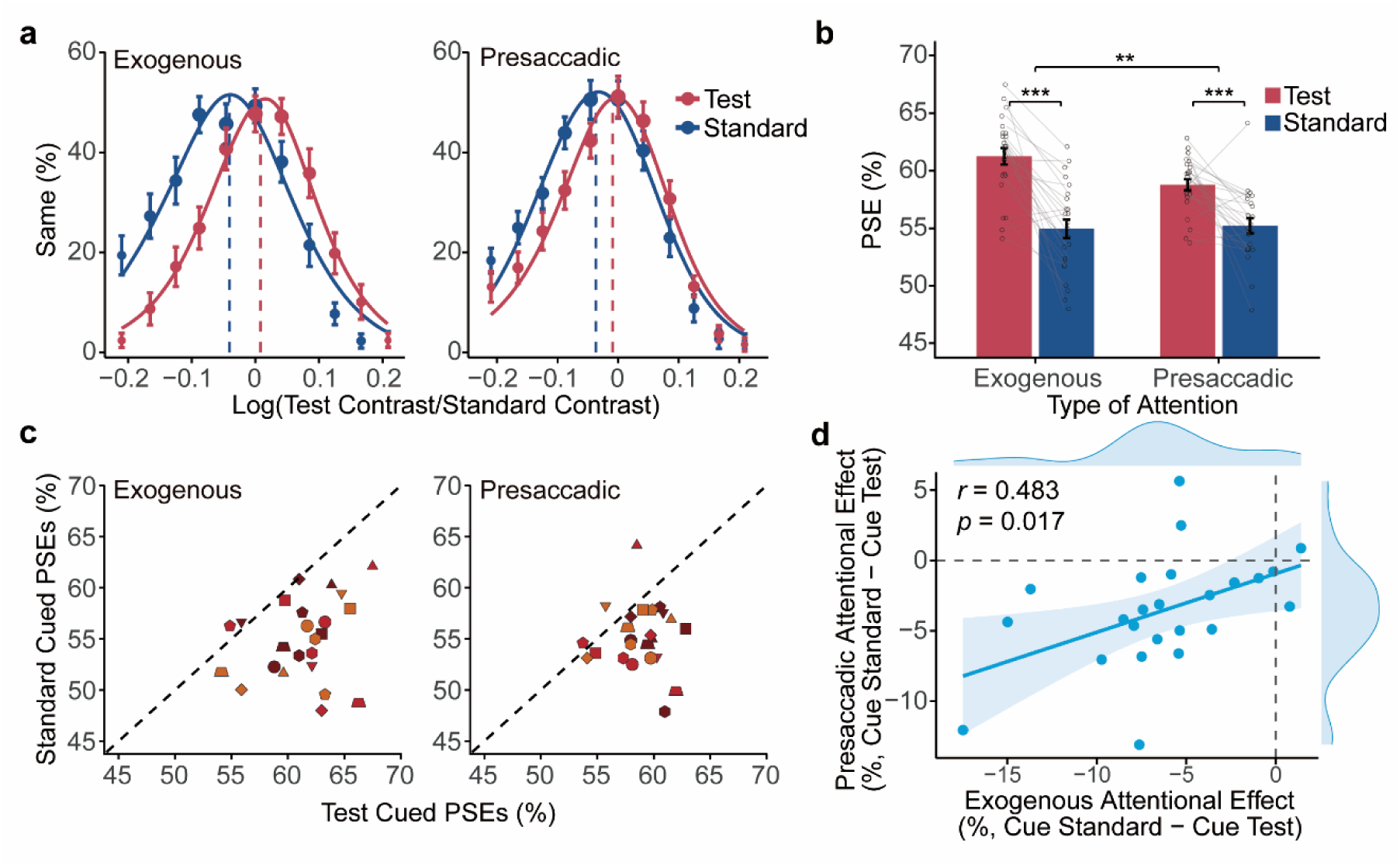
Results of Experiment 2. Psychometric functions in (a) show the mean percentage of ‘same’ responses relative to test stimuli contrast levels for the exogenous and presaccadic attention. The size of each data point is proportional to the number of trials corresponding to the contrast levels of the test grating. Panel (b) summarizes the averaged PSEs across cue conditions for both attention types, with the same participants’ data points connected. Individual PSEs for the standard-cued versus test-cued conditions are displayed in (c), with each participant identified by distinct shapes and colors. The diagonal dashed lines denote equal PSEs under test-cued and standard-cued conditions, suggesting no attentional effect on contrast appearance. Panel (d) shows the correlation between the effects of exogenous and presaccadic attention, with the solid blue line representing the linear regression line and shaded areas showing 95% confidence intervals. Marginal distributions reflect density distribution of attentional effects. Error bars represent ±1 standard errors of the mean. **p < .01. ***p < .001.

#### Correlation between attentional attenuation effects

The main purpose of Experiment 2 was to explore the potential association between voluntary presaccadic and involuntary exogenous attention. If a shared mechanism was involved, the two types of attention should produce significant correlated effects on contrast perception. As shown in Fig. 3d, there was a positive correlation between the effects of presaccadic and exogenous attention (*r* = .483, *p* = .017), indicating that participants who experienced a more intense attenuation effect with exogenous attention also had a more intense attenuation effect with presaccadic attention. This positive correlation suggests that presaccadic and exogenous attention might engage overlapping mechanisms in shaping subjective contrast perception, despite being allocated through different behavioral manners.

#### Temporal evolution of the presaccadic attentional effect

An important characteristic of presaccadic attention is that attentional effect gradually builds up during saccade preparation, peaking just before the saccade onset (H.-H. Li, Hanning, et al., 2021). If the attenuation effect is due to presaccadic attentional shift, we would expect to observe a temporal evolution of the attenuation effect as saccade onset approached. Our results indeed demonstrated a temporal profile like this (Fig. 4a and 4c). While the attentional attenuation effect was not significant at the early stage (122±31 ms before saccade onset) of saccade preparation (Fig. 4a; *t*(23) = 0.53, *p =* .60, Cohen’s *d* = 0.11, 95% CI for the mean difference = [−2.72, 1.61]), it became significant at the late stage (49±18 ms before saccade onset) of saccade preparation (*t*(23) = 5.72, *p* < .001, Cohen’s *d* = 1.17, 95% CI for the mean difference = [−6.00, −2.82]). The sliding window analysis further revealed a gradual build-up of attentional effect during saccade preparation, with significant attentional attenuation of contrast appearance emerging approximately 80 ms before saccade onset and peaking just before the onset of eye movement (Fig. 4c). This pattern is consistent with previous research on presaccadic attentional enhancement effect of contrast appearance for low-contrast stimuli (Rolfs & Carrasco, 2012), underscoring the progressive accumulation of attention during the preparation of saccadic eye movements.

**Fig. 4.**
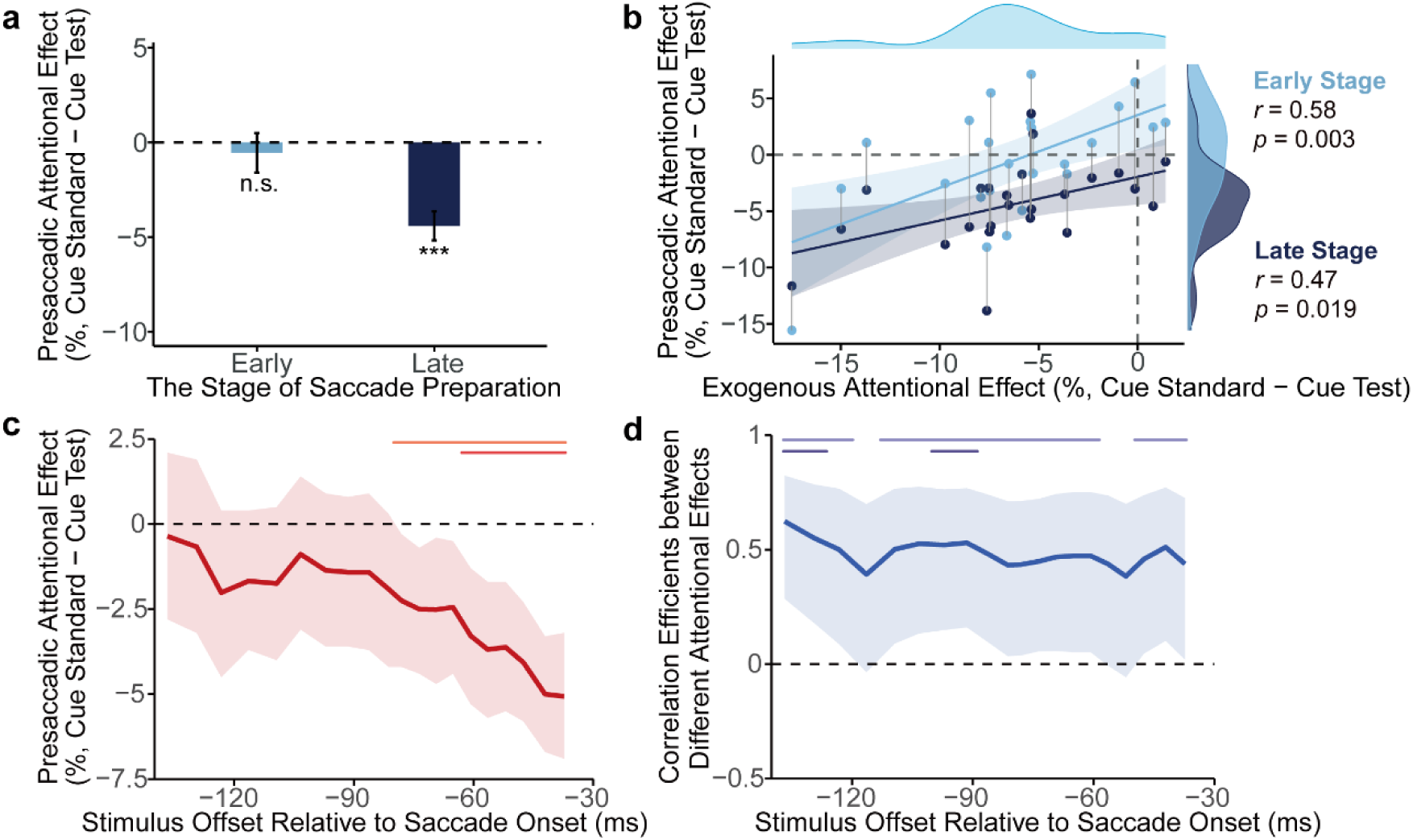
Temporal profile of the presaccadic attentional effect and its correlation with covert exogenous attention. Panel (a) displays the averaged presaccadic effects at early (distant from saccade onset) and late (proximal to saccade onset) saccade preparation stages. The error bars represent ±1 standard errors of the mean. ***p < .001. Panel (b) presents correlations of presaccadic attentional effect at early and late stages with exogenous attentional effect, where data for the same participants are connected with grey lines. The solid blue lines represent the linear regression lines with shaded areas for 95% confidence intervals. Marginal plots show the density distributions of presaccadic effects at both stages, as well as the exogenous attentional effect (identical to that in Fig. 3d). Panel (c) and (d) illustrate the sliding window analysis for the presaccadic attentional effect and the correlation efficient, respectively. The shaded areas represent the 95% confidence intervals. The upper horizontal lines indicate the significance intervals, with pink and light blue lines denoting periods of p < .05, and dark red and dark blue lines denoting periods of p < .01.

More importantly, we examined the temporal evolution of the correlation between presaccadic and exogenous attention (Fig. 4b and 4d). Intriguingly, positive correlations emerged at both the early and late stages of saccade preparation (Fig. 4b; early: *r* = 0.58, *p* = .003; late: *r* = 0.47, *p* = .019), unlike the temporal profile of the population-level attentional effect which reached significance only at the late stage. The sliding window analysis further revealed significant correlations throughout nearly the entire saccade preparation (Fig 4d). This persistent correlation throughout saccade preparation indicates that a qualitatively uniform mechanism drives the quantitative accumulation of attentional influence on visual perception, implying that a shared mechanism between presaccadic and exogenous attention is rapidly engaged from the initial moment of making a saccade.

## General Discussion

Whether attention alters our subjective visual experience of the world has been a topic of interest since the foundation of experimental psychology (Helmholtz Hv, 1896; James, 1890; Wundt, 1897). Our study contributes to this century-old inquiry by focusing on presaccadic attention, a form of attention ubiquitous in daily life. Understanding its effect on contrast perception, an elementary visual dimension, is crucial for comprehending the phenomenological correlates of attention in natural visual scenes. In Experiment 1, we found that presaccadic attention enhanced the appearance of low-contrast stimuli while attenuating the appearance of high-contrast stimuli, demonstrating a bi-directional modulation on contrast perception. This pattern is highly similar to that of covert exogenous attention but distinct from covert endogenous attention. In Experiment 2, we uncovered a positive correlation between the attentional effects of presaccadic attention and exogenous attention. This correlation emerged early and persisted just prior to the onset of a saccade. The similarity and correlation between the effects of voluntary presaccadic attention and involuntary exogenous attention suggest that shared mechanisms underlie these two types of attention.

The relationship between eye movement and covert attention is a long-standing debate. Whether covert exogenous and/or endogenous attention requires saccade preparation, as posited by the premotor theory of attention, is still contentious (Smith & Schenk, 2012). The conservative version of this theory argues that only covert exogenous attention depends on saccade preparation but covert endogenous attention operates independently of saccade preparation. Previous studies favoring (Casteau & Smith, 2020; Smith et al., 2004, 2012) or arguing against (Belopolsky & Theeuwes, 2009; Hunt & Kingstone, 2003; MacLean et al., 2015) this conserved version have primarily used accuracy or reaction time as measures (but see Hanning et al., 2019), which may conflate perceptual and non-perceptual factors and lead to the mixed results. Our results advance this debate by providing pure perceptual-level evidence: exogenous attention is closely associated with saccade preparation given their similar and correlated bi-directional attentional modulation of contrast appearance (Pan et al., 2023; Pan & Cai, 2022; Zhou et al., 2018), while endogenous attention should be independent of saccade preparation as it uniformly strengthens contrast appearance regardless of contrast levels (Luo et al., 2024).

The bi-directional modulation of contrast appearance by attention, though counterintuitive, might offer some behavioral benefits. Presaccadic attention is believed to facilitate peripheral information processing, and by amplifying the contrast appearance in low-contrast context, it makes faint peripheral details more discernible (Rolfs & Carrasco, 2012). However, uniform contrast enhancement may not be necessary for trans-saccadic vision. When a saccade target is briefly blanked during the perisaccadic period, visual perception of the saccade target improves, including location judgment (Deubel et al., 1996) and spatial frequency discrimination (Weiß et al., 2015), a phenomenon known as the “blanking effect”. Notably, the blanking effect occurs only for high-contrast targets (Matsumiya et al., 2016). The transient attenuation of high-contrast stimuli’s appearance by presaccadic attention is similar to brief visual blanking of a target, potentially facilitating peripheral information processing in high-contrast context. Combining our findings with those of others, we speculate that the blanking effect and presaccadic attentional effect might reflect common underlying mechanisms that support trans-saccadic vision, possibly also overlapping with exogenous attention processes.

The behavioral similarity between presaccadic and exogenous attention may also provide some inspirations and future directions for neural mechanism underlying eye movement and attention. Covert exogenous attention, a bottom-up process, is thought to operate primarily through feedforward mechanisms (Anton-Erxleben & Carrasco, 2013), and its attenuation effect on contrast appearance is indeed mediated by modulation of initial afferent V1 activity (Pan et al., 2023). In contrast, presaccadic attention is hypothesized to be driven by feedback signals from higher-order cortex responsible for controlling saccades, such as the FEF and IPS (Moore & Zirnsak, 2017), which project toward the visual cortex (Rolfs, 2015). How do these two distinct forms of attention similarly modulate early visual cortex activity? The superior colliculus (SC), a crucial structure for both overt and covert attention shifting (Krauzlis et al., 2013), is a likely candidate. SC neurons in the intermediate layer are modulated by saccade preparation and exogenous cues, but not by endogenous cues (Ignashchenkova et al., 2004), providing a neural basis for the behavioral differences between presaccadic, exogenous, and endogenous attention. Additionally, SC plays an important role in relaying feedback signals from oculomotor areas like the FEF to the visual cortex (Hu et al., 2019). Recent evidence further indicated that SC not only receives input from V1 but encodes visual saliency earlier than V1 (White et al., 2017) and even modulates activity of V1 as well as LGN (Ahmadlou et al., 2018), suggesting its potential role in attentional modulation of sensory cortex. Future electrophysiological and neuroimaging studies can investigate how these regions interact to drive attentional modulation of sensory perception.

In conclusion, our findings establish a strong connection between voluntary presaccadic attention and covert exogenous (involuntary) attention as evidenced by their similar and correlated perceptual consequences. This suggests that attention shifts preceding goal-directed eye movements differ qualitatively from endogenous attention that occur when the eyes remain stationary, and, more unexpectedly, resemble exogenous attention in the absence of eye movements. Our findings point to a shared attentional modulation mechanism that underlies both voluntary presaccadic and involuntary transient attention. These findings provide new insights into neural and computational research concerning the function of eye movement and attention in various scenarios of visual perception.

## Transparency

## Author Contributions

**Tianyu Zhang:** Conceptualization; Data curation; Formal analysis; Investigation; Methodology; Software; Visualization; Writing – original draft.

**Yongchun Cai:** Conceptualization; Data curation; Formal analysis; Funding acquisition; Investigation; Methodology; Project administration; Supervision; Writing – review & editing.

## Open Practices

No aspects of the study were preregistered. All data and analytic codes can be made available upon request to the first author.

